# Dynactin has two antagonistic regulatory domains and exerts opposing effects on dynein motility

**DOI:** 10.1101/110775

**Authors:** Takuya Kobayashi, Takuya Miyashita, Takashi Murayama, Yoko Y. Toyoshima

## Abstract

Dynactin is a dynein-regulating protein that increases the processivity of dynein movement on microtubules. Recent studies have shown that a tripartite complex of dynein–dynactin–Bicaudal D2 is essential for highly processive movement. To elucidate the regulation of dynein motility by dynactin, we focused on two isoforms (A and B) of dynactin 1 (DCTN1), the largest subunit of dynactin that contains both microtubule- and dynein-binding domains. The only difference between the primary structures of the two isoforms is that DCTN1B lacks the K-rich domain, a cluster of basic residues. We measured dynein motility by single molecule observation of recombinant dynein and dynactin. Whereas the tripartite complex containing DCTN1A exhibited highly processive movement, the complex containing DCTN1B dissociated from microtubules with no apparent processive movement. This inhibitory effect of DCTN1B was caused by reductions of the microtubule-binding affinities of both dynein and dynactin, which is attributed to the coiled-coil 1 domain of DCTN1. In DCTN1A, the K-rich domain antagonized these inhibitory effects. Therefore, dynactin has two antagonistic domains and promotes or suppresses dynein motility to accomplish correct localization and functions of dynein within a cell.

## Introduction

Dynactin is a very large multi-subunit complex that links dynein with its cargo and is known as a cytoplasmic dynein modulator by binding dynein to specific vesicles or organelles [1–3]. Cytoplasmic dynein is a minus end-directed multi-subunit microtubule motor protein [4]. Dynactin is involved in many cellular functions, including vesicle transport [2,5], organelle positioning [6,7], spindle assembly [8] and microtubule plus end localization [9–11] with dynein. Dynactin abnormalities cause several diseases, including Perry syndrome [12,13] and amyotrophic lateral sclerosis [14,15]. Dynein–dynactin (DD) complexes are distributed broadly range in cells and their spatial distribution changes throughout the cell cycle [16]. Correct localization of the DD complex is important for maintenance of cellular functions. However, the regulatory mechanism of the DD complex in determining its cellular distribution has been poorly understood.

Previous studies have indicated that dynactin enhances microtubule binding of dynein and induces processive movement of dynein using beads coated with multiple dynein molecules [17,18]. More recently, it has been reported that the DD complex itself does not exhibit unidirectional and highly processive movements. However, the dynein–dynactin–Bicaudal D2 (BICD2) (DDB) complex has exhibited unidirectional and highly processive movements in single molecule observations [19,20].

Dynactin itself is a very large multi-subunit complex including dynactin 1 (DCTN1) [21], p50, actin-related protein 1 (Arp1) [22], and other eight kinds of subunits [1]. DCTN1, also known as p150^Glued^, forms a dimer and extrudes the Arp rod as the arm and shoulder, containing a microtubule-binding region (N-terminal region) [23], dynein-binding region [coiled-coil 1 (CC1) domain] [24], and Arp1-binding region (C-terminal region) [3]. The N-terminal region consists of a CapGly domain [25,26] and K-rich domain [27]. The CapGly domain exhibits equimolar binding to microtubules [28], and the K-rich domain is involved in supporting this interaction [28], which is similar to the K-loop of kinesin [29,30]. The N-terminal region of DCTN1 is different between spliced isoforms, DCTN1A (1A isoform) and DCTN1B (1B isoform). The 1A isoform contains both CapGly and K-rich domains, whereas the 1B isoform lacks the K-rich domain [27]. These isoforms are differentially expressed in tissues [27,31]. However, the roles of these isoforms have not been elucidated in DD and DDB complexes.

In this study, we used recombinant DCTN1 to isolate the dynein complex with one DCTN1 isoform and conducted single molecule observations to quantify the microtubule-binding ratio of dynein and dynactin. Consequently, we demonstrate that the two DCTN1 isoforms (1A and 1B) exert different effects on dynein motility. The 1A isoform induces unidirectional movement, whereas the 1B isoform reduces the microtubule-binding affinity to inhibit unidirectional movement. By comparing the properties of the two isoforms and several mutants, we show that both CapGly and K-rich domains are essential for microtubule binding of dynein to promote unidirectional movement. Furthermore, we found that the CC1 domain is responsible for reductions of the microtubule-binding affinities of both dynein and dynactin and that the K-rich domain antagonizes these effects. We conclude that DCTN1 has two antagonistic regulatory domains that interact intramolecularly with each other and then exerts opposing effects on dynein motility.

## Materials and Methods

### Construction of expression vectors

cDNAs encoding DCTN1 isoforms and BICD2 were amplified from HEK293 cells by RT-PCR. The PCR primers used for cloning DCTN1 isoforms were as follows: 5′-ATGGCACAGAGCAAGAGGCAC-3′ and 5′-TTAGGAGATGAGGCGACTGTG-3′ for DCTN1 (NM_004082), and 5′-ATGTCGGCGCCGTCGGAGGAG-3′ and 5′-CTACAGGCTCGGTGTGGCTGGCTTGG-3′ for BICD2 (NM_015250). Note that our cloned DCTN1 sample from HEK293 cells was 1B isoform which lacked 21 amino acids of the K-rich domain (∆exon 5–7) as reported by Dixit et al [32]. cDNA for the 1A isoform was constructed by inverse PCR using the following PCR primers: 5′-GCCCGAAAGACCACAACTCGGCGACCCAAGCCCACGCGCCCAGCCAGTACT-3′ and

5′-TGTCGGTGCCTTCTTAGGCTTCAGTCCCCGCAGTTTGCTAGTCTTTGCAGT-3'. For 1B∆CC1 mutant construction, the CC1 domain-coding region was deleted using the following PCR primers: 5′-CCACCTCCAGAGACCTTTGAC-3′ and 5′-TGGGGAAGGAAGCGGGGGGAC-3′.

The PCR products were cloned into pcDNA5/FRT/TO (LifeTechnologies), a tetracycline-inducible mammalian expression vector. For protein purification, the streptavidin-binding peptide (SBP)-tag-inserted multifunctional green fluorescent protein (GFP)-tag [33] was fused to the N-terminus of DCTN1. The insertion of these tags or deletion of specific domains was achieved by inverse PCR.

### Generation of stable inducible HEK293 cell lines

Flp-In T-REx HEK293 cells (Life Technologies, Carlsbad, CA) were maintained in Dulbecco’s modified Eagle’s medium (Nacalai, Tokyo, Japan) supplemented with 10% fetal bovine serum and 2 mM L-glutamine. Stable inducible cell lines were generated by co-transfection of the pcDNA5/FRT/TO vector containing recombinant DCTN1 isoforms or their mutants with the pOG44 vector encoding Flp recombinase according to the manufacturer’s instructions. The transfectants were screened by selection with 100 μg/ml hygromycin, followed by harvesting hygromycin-resistant colonies.

### Protein purification

HEK293 cells expressing DCTN1 isoforms, dynein heavy chain, or BICD2 were cultured in 20 150-mm culture dishes, and then collected and rinsed twice with phosphate-buffered saline. Cell lysates were prepared by homogenizing the cells in buffer B (10 mM Pipes-NaOH, pH 7.0, 200 mM NaCl, 10% sucrose, and 1 mM dithiothreitol) containing 0.05% Triton X-100 and complete mini protease inhibitor cocktail (Roche, Basel, Swiss). After centrifugation and filtration, the lysates were applied to a StrepTrap HP column (GE healthcare, Amersham, UK) that had been equilibrated with buffer B. After washing with buffer A (10 mM Pipes-NaOH, pH 7.0, 150 mM NaCl, and 1 mM dithiothreitol), bound proteins were eluted with buffer B containing 2.5 mM desthiobiotin. Purified DCTN1 isoforms and mutants were confirmed by sodium dodecyl sulfate-polyacrylamide gel electrophoresis (SDS-PAGE) (S1 Fig).

The CC1 fragment consisted of 214–547 a.a. of p150^Glued^. The 1BN^-^GCN4 fragment consisted of 1–193 a.a. of the 1B isoform and the GCN4 sequence [34] to dimerize this fragment. These fragments were fused with GFP and a His-tag at the C-terminus. CC1 and 1BN-GCN4 fragments were expressed in pET and pCold expression systems, respectively. These fragments were then purified using profinity IMAC Ni-charged Resin (Bio-Rad, Hercules, CA).

Tubulin was purified from porcin brain as described by Weingarten [35]. 134 Tubulin was incubated for 30 min at 37°C with 1 mM GTP and 5 mM MgSO_4_ to 135 polymerize. After the incubation, taxol was added to a final concentration of 40 μM.

### Single molecule observation

Flow chambers were constructed with silanized glass, which were coated with a 5% anti-β tubulin antibody solution and then blocked with 10% Pluronic F-127 and casein. Taxol-stabilized microtubules were introduced into the flow chamber to obtain a constant density of microtubules on the glass surface in each experiment, followed by Alexa 647-labeled dynein and GFP-labeled dynactin in SM buffer (12 mM Pipes-KOH, pH 6.8, 25 mM KAce, 1 mM ATP, 2 mM DTT, 10% saturated casein, and an oxygen-scavenging system). Single molecule observations were performed at 25°C using a total internal reflection fluorescence (TIRF) microscope (IX71, Olympus, Tokyo, Japan) equipped with a 100×/1.45 PlanApo objective lens (Olympus). Images were acquired with a back illumination EMCCD camera (iXon, DV887DCS-BV, Andor, UK) at an exposure time of 100 ms.

Dynein and dynactin behaviors were analyzed by recording fluorescent spots that were visible along microtubules for more than 1 s. The numbers of dynein and dynactin molecules were counted per microtubule length of 10 μm for 10 s under the condition of 1 nM dynein or dynactin molecules. The attachment rate was the number of newly detected molecules on a microtubule for 10 s. Spots of dynein that moved bidirectionally for more than 400 nm were classified as “diffusive”, those that moved bidirectionally for less than 400 nm were classified as “stationary”, and those that moved to the minus end by more than 400 nm were classified as “unidirectional”.

### Protein binding assay

Dynein and dynactin were mixed in assay buffer (10 mM Pipes-NaOH, pH 7.0, 75 mM NaCl, 45 mM imidazole, and 0.01% Tween 20), and then the mixture was incubated for 10 min at 25°C. TALON Dynabeads (Life Technologies, Carlsbad, CA) were added to the mixture, followed by incubation for 10 min at 25°C. The Dynabeads were collected by a magnet. Unbound protein was recovered and washed twice with assay buffer. The bound protein was eluted by elution buffer (10 mM Pipes-NaOH, pH 7.0, 150 mM NaCl, and 300 mM imidazole). Unbound and bound fractions were analyzed by SDS-PAGE.

## Results

### Motility of the DDB complex containing 1A or 1B isoforms

Previous studies have indicated that the DDB complex exhibits highly processive and unidirectional movements [19,20]. However, the difference in DCTN1 isoforms of dynactin remains unknown in the DDB complex. To investigate whether DCTN1 isoforms influence the highly processive movement of DDB complexes, we observed the behavior of DDB complexes with different DCTN1 isoforms (1A and 1B) by TIRF microscopy (Fig 1). To this end, each DCTN1 isoform was fused with a multifunctional GFP-tag at the N-terminus and expressed in HEK293 cells (Fig 1a). These DCTN1 isoforms were incorporated into the endogenous dynactin complex and successfully purified by affinity chromatography (S1 Fig).

**Fig 1.**
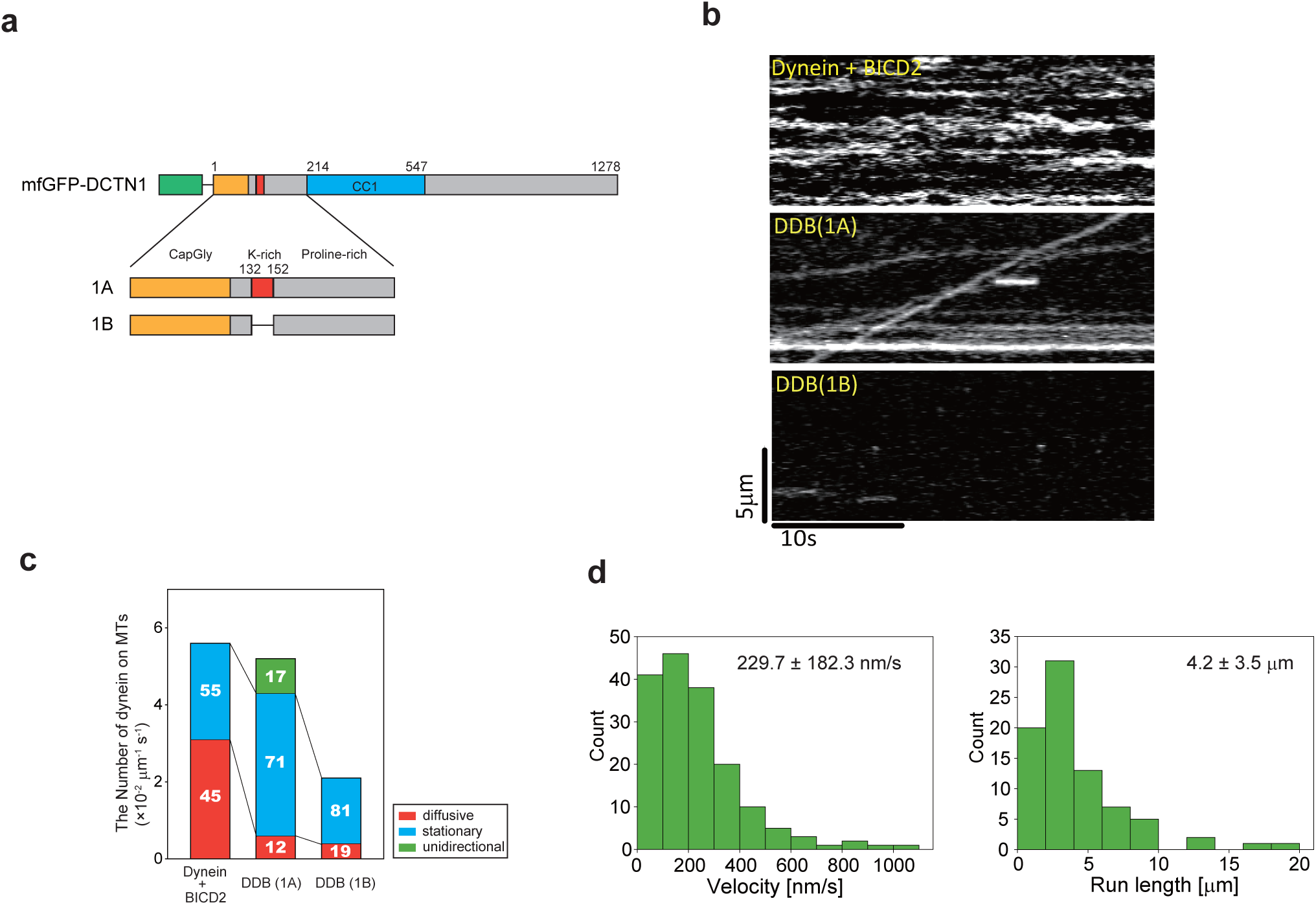
Behaviors of DDB complexes with different DCTN1 isoforms. (a) N-terminal regions of DCTN1 isoforms. Amino acid numbering is based on p150^glued^. (b) Kymographs of dynein movement on microtubules (MTs) in the presence of 1 nM Alexa 647-labeled dynein, 2 nM 1A or 1B isoforms, and 20 nM BICD2. (c) Quantification of the behavior of dynein molecules on MTs. The quantification data are presented as segmented vertical bars: the number of diffusive (red), stationary (blue), or unidirectional (green) dynein molecules on MTs. Quantification of dynein molecules on MTs in the presence of each dynactin isoform or mutant. The number of dynein molecules on MTs in a time window (10 s) was normalized to the microtubule length (10 μm) under the condition of 1 nM Alexa 647-labeled dynein and 2 nM 1A or 1B isoforms. Mean ± S.D., *n* = 9 windows. \**P<* 0.01, Student’s *t*-test. (d) Histograms of the velocity and run length of unidirectional movement of the DDB complex (1A). Mean ± S.D., *n* = 168 particles (velocity) and *n* =80 particles (run length).

We observed the behavior of Alexa647-labeled dynein on microtubules (Fig 1b) and classified it into three types, i.e., diffusive, stationary and unidirectional movement (Fig 1c). The number of dynein molecules exhibiting each type of movement was calculated based on our counting criteria (see Materials and Methods). In the absence of dynactin (dynein + BICD2), the number of dynein molecules on microtubules was 5.6 ± 0.8. Dynein was either stationary (55%) or diffusive (45%) and no processive movement was observed (Fig 1b, 1c and S1 Table). Addition of the 1A isoform (DDB 1A) did not change the number of dynein molecules on microtubules (5.2 ± 1.0). However, a significant fraction of dynein (17%) moved unidirectionally with marked reduction in the diffusive fraction (12%) and an increase in the stationary fraction (71%) compared with the absence of dynactin. The unidirectional movement occurred with a mean velocity of 229.7 ± 182.3 nm/s and a mean run length of 4.2 ± 3.5 μm (Fig 1d). The highly processive movement of the DDB complex with the 1A isoform is consistent with previous reports [19,20].

In contrast, addition of the 1B isoform (DDB 1B) resulted in a significantly lower number of dynein molecules on microtubules (2.1 ± 0.7, *P* < 0.01) than without dynactin or with the 1A isoform. Most dynein was stationary (81%) and the remaining fraction was diffusive (19%). No unidirectional movement was observed (Figs 1b and 1c). These results suggest that the 1A isoform is essential for unidirectional processive movement of the DDB complex. Furthermore, the fact that addition of the 1B isoform reduced the number of dynein molecules on microtubules suggests that the 1B isoform may inhibit microtubule binding of dynein in the DDB complex.

### Role of the DCTN1 isoform in the behavior of DD complexes on microtubules

To investigate the effect of DCTN1 isoforms on dynein motility in a more simple system, we observed the behavior of DD complexes on microtubules in the absence of BICD2 (Fig 2a). The number of dynein molecules on microtubules was 7.1 ± 1.7 molecules in the absence of dynactin (Fig 2b). Seventy-six percent of the observed dynein exhibited diffusive motion and 24% of dynein molecules were stationary (Fig 2b and S1 Table). The number of dynein molecules on microtubules was slightly decreased in the presence of the 1A isoform (5.6 ± 1.0 dynein molecules, *P* < 0.01). The fraction of stationary dynein molecules was increased to 59%, and 27% of dynein molecules exhibited diffusive motion. Surprisingly, 14% of dynein molecules exhibited unidirectional processive movement (S1 Movie). The mean velocity and run length of the unidirectional movement were 130.0 ± 79.5 nm/s and 1.6 ± 0.9 μm, respectively (Fig 2c), which were lower than those of the DDB complex (see Fig 1d). Conversely, in the presence of the 1B isoform, the number of dynein molecules was drastically decreased to 0.6 ± 0.5 (Figs 2a and 2b). Whereas 79% of dynein in the presence of the 1A isoform did not dissociate from microtubules during the observation (30 sec) (Fig 2a), the duration of dynein molecules on microtubules was greatly reduced in the presence of the 1B isoform (τ = 1.2 s) compared with the 1A isoform (Fig 2d). These findings indicate that the 1A isoform is necessary and sufficient for the unidirectional movement, and that the 1B isoform inhibits the microtubule binding ability of dynein.

**Fig 2.**
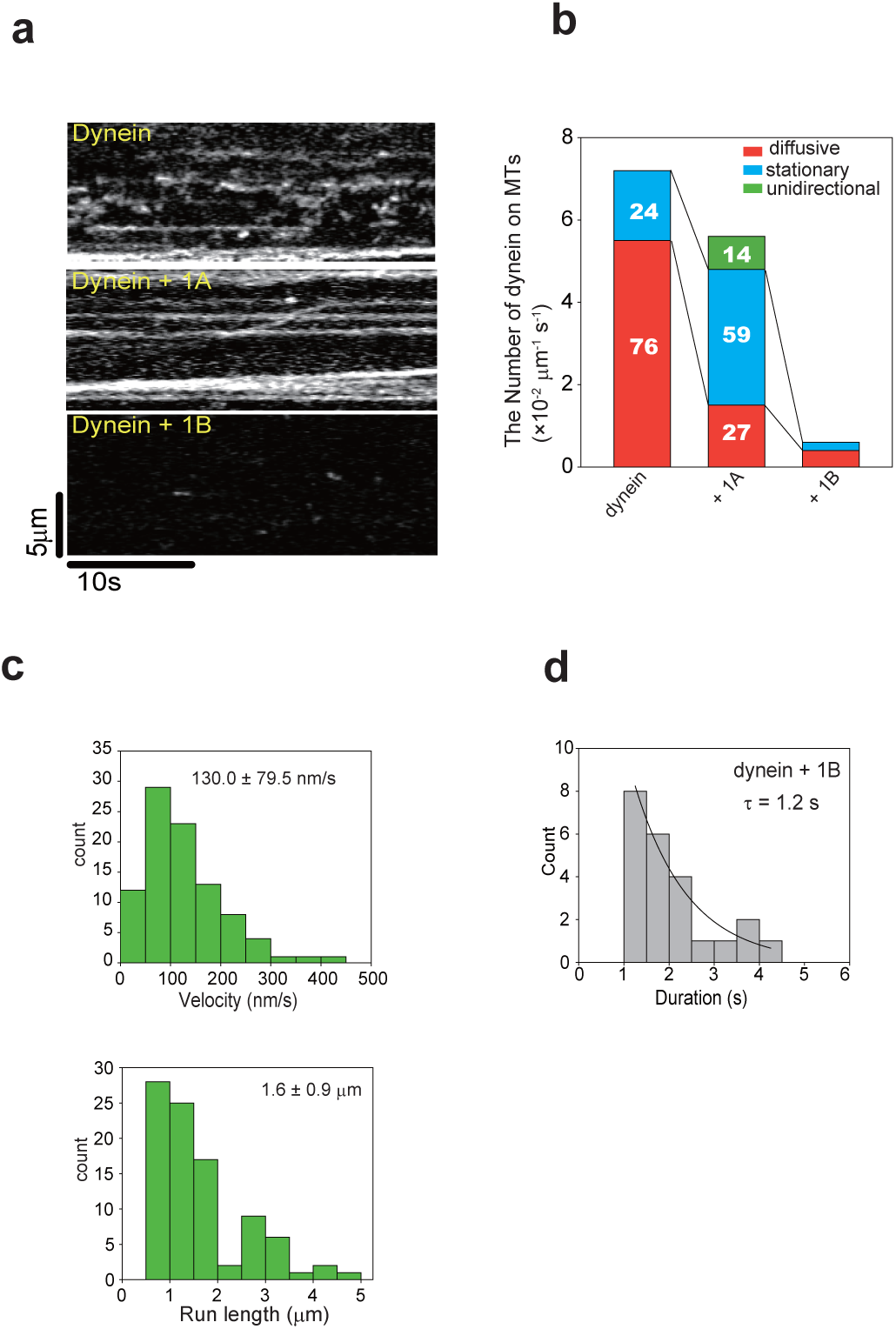
Behaviors of DD complexes with different DCTN1 isoforms. (a) Kymographs of dynein movement on microtubules (MTs) in the presence of each dynactin isoform. (b) The number of dynein molecules on MTs was the same as that in Fig. 1c under the condition of 1 nM Alexa 647-labeled dynein and 2 nM of each dynactin isoform. Mean ± S.D., *n* = 18 windows (dynein alone), *n* = 15 windows in the presence of 1A or 1B isoforms. The number of dynein molecules under each condition was altered significantly compared with dynein alone (*P* < 0.01, Student’s *t*-test). Quantification of the behavior of dynein molecules on MTs is presented as segmented vertical bars: the number of diffusive (red), stationary (blue), or unidirectional (green) dynein molecules on MTs. The number of dynein molecules on MTs was the same as that in Fig 1b. Mean ± S.D. (c) Histograms of the velocity and run length of unidirectional movement of dynein molecules in the presence of the 1A isoform. *n* = 92 particles. (d) Duration of dynein interacting with MTs in the presence of 1B. *n* = 23 particles (1B isoform).

### The CC1 domain inhibits dynein motility

The above results clearly demonstrated that the 1B isoform inhibits the microtubule-binding ability of dynein in both the DDB complex (Fig 1) and DD complex (Fig 2). We hypothesized that the binding of dynactin with the 1B isoform to dynein affects its microtubule-binding ability. It is known that the CC1 domain of DCTN1 binds to the dynein intermediate chain [24], and the Arp1-rod interacts with the dynein tail [36]. Therefore, we investigated the effect of the CC1 domain on the microtubule binding ability of dynein.

We constructed the CC1 fragment and the 1B isoform lacking the CC1 domain (1BΔCC1) (Fig 3a) and initially examined the binding ability of the CC1 domain to dynein by pull-down assays with the purified proteins (Figs 3b and 3c). The 1B isoform bound to dynein in a simple 1:1 binding ratio with a dissociation constant (kd) of 2.3 nM. The engineered CC1 fragment also exhibited a similar binding ability with a dissociation constant (k_d_) of 2.0 nM. Conversely, the 1B isoform lacking the CC1 domain (1B∆CC1) did not specifically bind to dynein. These results indicate that the CC1 domain is the primary dynein-binding site of dynactin.

**Fig 3.**
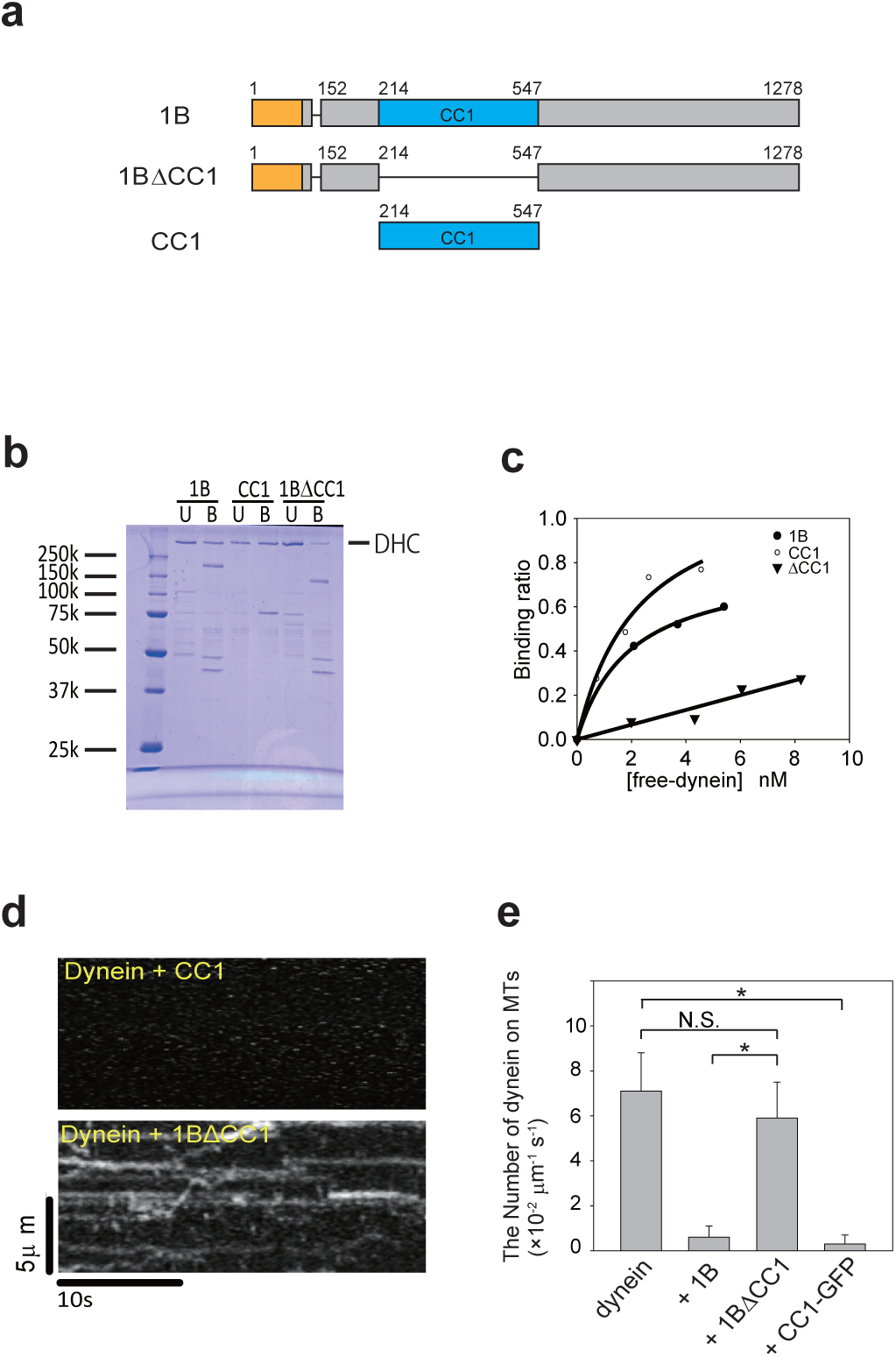
The CC1 fragment binds to dynein and inhibits microtubule binding of dynein. (a) Mutant of the 1B isoform and the CC1 fragment. Amino acid numbering is based on p150^glued^. (b) Interactions between dynein and dynactin determined by TALON Dynabeads pull-down assays of purified proteins. The protein bands of dynein heavy chain (DHC) in SDS-PAGE gels were quantified by densitometry, (c) Binding ratio of dynein with the 1B isoform (○), CC1 fragment (•), and 1BΔCC1 mutant (▾). (d) Kymographs of dynein motility on microtubules (MTs) in the presence of the 1B∆CC1 mutant or CC1 fragment. (e) Quantification of dynein molecules on MTs in the presence of the 1B∆CC1 mutant or CC1 fragment. The number of dynein molecules on MTs was the same as that in Fig 1b under the condition of 1 nM Alexa 647-labeled dynein and 2 nM 1B∆CC1 mutant or CC1 fragment. Mean ± S.D., *n* = 15 windows. \**P* < 0.01, Student’s *t*-test. N.S., not significant.

We next examined the effect of the CC1 fragment and 1B∆CC1 mutant on the microtubule-binding ability of dynein (Fig 3d). The CC1 fragment greatly reduced the number of dynein molecules (0.3 ± 0.4, Fig 3e) to a level similar to that with the 1B isoform (0.6 ± 0.5, Fig 2b). In contrast, the number of dynein molecules in the presence of the 1B∆CC1 mutant did not significantly differ (5.9 ± 1.6, Fig 3e) compared with dynein alone (7.1 ± 1.7, Fig 2b). These results demonstrate that the CC1 domain binds to dynein and inhibits the microtubule-binding ability of dynein.

### Microtubule-binding affinities of DCTN1 isoforms

The fact that the CC1 domain binds to dynein and inhibits its microtubule binding in the 1B isoform raises an important issue: why the 1A isoform does not inhibit the microtubule binding of dynein and enables unidirectional processive movement. Dynactin itself interacts with microtubules via the N-terminal region of DCTN1 that contains a CapGly domain [23,28]. Because the K-rich domain is reported to increase the microtubule-binding ability of the CapGly domain, we hypothesized that the microtubule-binding ability of each DCTN1 isoform might be different. To estimate the microtubule-binding abilities of DCTN1 isoforms, we observed the behavior of dynactin on microtubules by TIRF microscopy. The 1A isoform was frequently observed on microtubules (Fig 4b) and the number of dynactin molecules on microtubules was 6.6 ± 1.3 (Fig 4c). In contrast, the 1B isoform exhibited a 10-fold reduction in the number of dynactin molecules on microtubules (0.6 ± 0.6).

**Fig 4.**
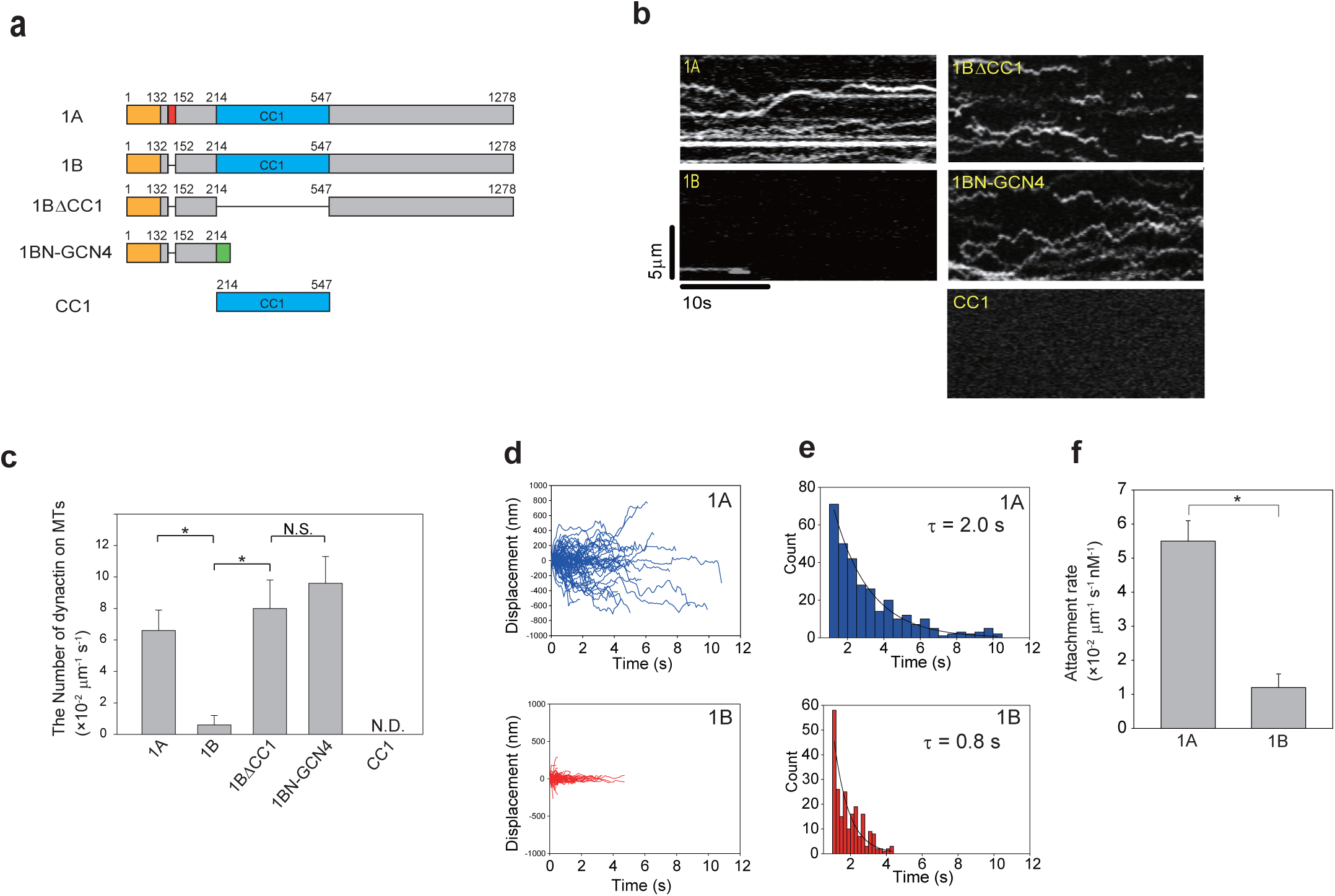
Single molecule behavior of dynactin on microtubules. (a) Mutant of the 1B isoform and the fragments of DCTN1. Amino acid numbering is based on p150^glued^. (b) Kymographs of dynactin including each isoform and mutant behavior on microtubules (MTs). (c) Quantification of each isoform and mutant on MTs. The number of dynactin molecules on MTs was the same as that in Fig 1b. Mean ± S.D., *n* = 9 windows (1A isoform, 1B∆CC1 mutant, 1BN-GCN4 fragment and CC1 fragment) and *n* = 6 windows (1B isoform). \**P*< 0.01, Student’s *t*-test. (d) Time course of displacement of 1A (upper panel) and 1B (lower panel) isoforms on MTs. The diffusion coefficients of 1A and 1B isoforms were 55.4 × 10^2^ and 1.9 × 10^2^ nm^2^/s, respectively. (e) Duration of 1A (upper panel) or 1B (lower panel) isoforms on MTs. *n* = 487 particles (1A isoform) and *n* = 454 particles (1B isoform). (f) Attachment rates of 1A and 1B isoforms. The attachment rate was normalized to the microtubule length (10 μm) and observation time (10 s) under the condition of 1 nM GFP-fused dynactin. Mean ± S.D., *n* = 30 microtubules (1A isoform), *n* = 28 microtubules (1B isoforms), *P < 0.01, Student’s *t*-test.

The 1A isoform exhibited highly diffusive movements with a diffusion coefficient of 46.9 × 10^2^ nm^2^/s (Fig 4d and S2 Fig) and interacted with microtubules for an average of 2.0 s (Fig 4e). Conversely, the 1B isoform moved much less diffusively with a diffusion coefficient of 2.0 × 10^2^ nm^2^/s (Fig 4d and S2 Fig) and the dwell time on microtubules was 0.8 s (Fig 4e). The attachment rates of 1A and 1B isoforms were 5.5 ± 0.6 and 1.2 ± 0.4, respectively (Fig 4f). Because the dissociation constant is proportional to the product of the attachment rate and duration, the dissociation constants of 1A and 1B isoforms were 0.1 × 10^2^ μm nM and 1.0 × 10^2^ μm nM, respectively. Thus, the microtubule-binding affinity of the 1A isoform was 10-fold more than that of the 1B isoform. The observed number of molecules on microtubules (see Fig 4c) was consistent with the microtubule-binding affinity, indicating that the number of molecules based on our criteria is a good measure of the microtubule-binding affinity.

These results suggest that the 1B isoform has much lower microtubule-binding ability than the 1A isoform. Thus, the K-rich domain represses the inhibitory effect of the CC1 domain on the microtubule-binding ability of the 1A isoform.

To examine the effect of the CC1 domain on the microtubule-binding ability of dynactin, we deleted the CC1 domain from the 1B isoform (1B∆CC1). Interestingly, the 1B∆CC1 mutant interacted with microtubules (8.0 ± 1.8), which was comparable to the 1A isoform (6.6 ± 1.3). Similarly, the 1BN-GCN4 fragment, which was the dimerized N-terminal fragment of the 1B isoform induced by the GCN4 sequence, interacted with microtubules at a similar level (9.6 ± 1.7, Fig 4c). The CC1 fragment itself did not interact with microtubules (Fig 4c). These results suggest that the N-terminal region is a unique site in the dynactin complex to interact with microtubules. The microtubule-binding ability of the CapGly domain is reduced by the CC1 domain and the K-rich domain antagonizes the inhibitory effect of the CC1 domain.

## Discussion

In this study, we found that dynactin has two agonistic regulatory domains (CC1 and K-rich domains) and exerts opposing effects on dynein motility depending on the DCTN1 isoform. The function of dynactin to increase dynein processivity, which has already been shown in previous reports [19,20], is considered to be attributed to the 1A isoform. Other than neurons, most tissues express more of the 1B isoform than the 1A isoform [27,31]. It appears that both 1A and 1B isoforms of DCTN1 coexisted in the previous reports [37,38], and the observed highly processive movement was achieved by the fraction with the 1A isoform. While the effect of the 1A isoform has been observed in previous studies, the properties of the 1B isoform were not revealed. In this study, we used recombinant DCTN1 isoforms and each isoform was investigated separately. Recombinant DCTN1 was incorporated into the dynactin complex and affinity purified using the SBP-tag in the fused multifunctional GFP [33] (S1 Fig). This purification method made it possible to isolate the dynactin complex with one DCTN1 isoform or mutant to compare the functions of the isoforms in dynein motility. Moreover, we evaluated the microtubule-binding ability of dynein by quantitative single molecule analysis. As a result, we were able to determine the unbinding property of DD complexes. So far, the unbinding property has not been assessed by single molecule analysis because it is an invisible phenomenon. Our strategies revealed new functions of several domains within DCTN1.

In the presence of the 1A isoform, we detected a decrease in the fraction of diffusive molecules of the DD complex, but an increase in that of stationary molecules compared with dynein alone (Fig 2b). The increase in stationary dynein might be due to the increase in microtubule-binding affinity, because the number of microtubule-binding sites in a DD complex is twice that in dynein alone. Alternatively, while diffusive motion does not appear to be involved in the unique binding of dynein to microtubules, dynactin might increase the unique binding of dynein, leading to the increase in stationary fractions. The DD complex with the 1A isoform generated unidirectional movement in a manner similar to that of multimolecular dyneins [39]. Because the DDB complex showed higher velocity and processivity (Fig 1d) than the DD complex (Fig 2c), BICD2 appears to regulate the complex to achieve a stable state that is favorable for the unidirectional movement and then increases processivity.

Conversely, the 1B isoform reduced the microtubule-binding affinity of dynein in both DDB and DD complexes (Figs 1c and 2b). Thus, the 1B isoform has an inhibitory effect on dynein motility. Because the only difference between the two isoforms is that the 1B isoform lacks the K-rich domain, this domain is important to support the microtubule-binding ability of the CapGly domain. Furthermore, the lack of the K-rich domain caused dissociation of the DDB complex (DDB 1B) (Fig 1c) and DD complex (Fig 2b). Therefore, the K-rich domain represses the dissociation of DDB and DD complexes by the 1B isoform (Fig 5, right, blue dashed line).

**Fig 5.**
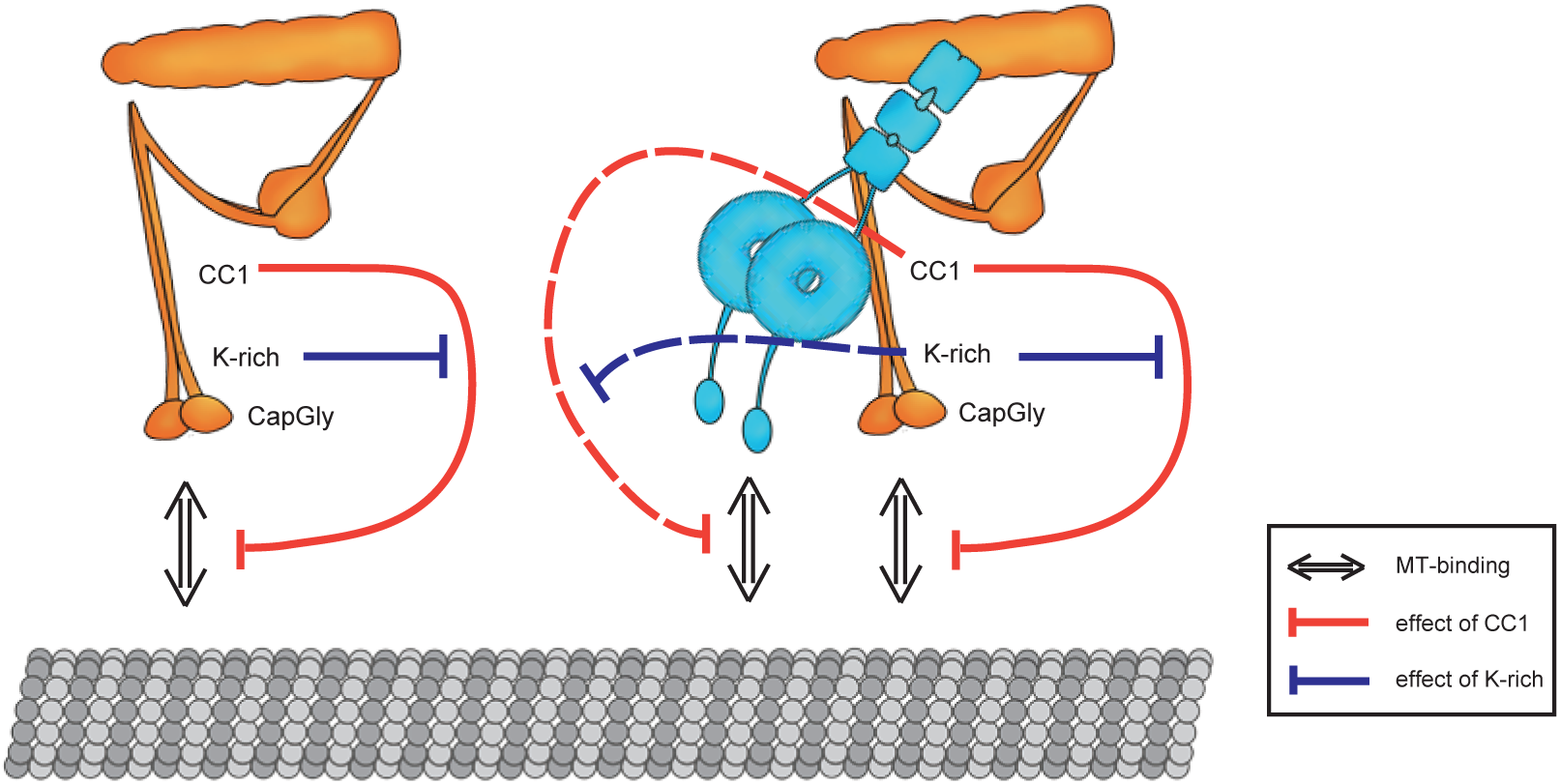
Relationships between regulatory domains. Left: The CC1 domain inhibits the microtubule-binding ability of the CapGly domain within the same molecule, and this inhibition is repressed by the K-rich domain. Right: The effect on dynactin itself still remains. The CC1 domain, which is bound to dynein, inhibits the microtubule-binding ability of the DD complex, and this inhibition is repressed by the K-rich domain.

The inhibition of dynein motility could thus be executed by non-N-terminal regions. Therefore, we focused on the CC1 domain that contains the dynein-binding site. The CC1 fragment, CC1-GFP, reduced the microtubule-binding affinity of dynein (Fig 3e), suggesting that the CC1 domain is a regulatory unit to inhibit dynein motility (Fig 5, right, red dashed line). This result is supported by a previous study using dynein molecule-coated beads [40]. Based on previous studies, the CC1 fragment is known to be a dynein inhibitor because the addition of excess CC1 fragments disrupts the DD complex by competitive inhibition with endogenous dynactin in cells [41,42]. Our finding of the inhibitory effect of CC1 on dynein motility *in vitro* suggests that the inhibition of dynein motility in a cell is not only caused by disruption of DD complex but also by the direct inhibitory effect of CC1.

The inhibitory mechanism of the microtubule-binding ability of dynein by the CC1 domain is unknown. The CC1 domain is reported to contain the primary dynein-binding site that binds to the dynein intermediate chain [24,43,44]. Because the dynein heavy chain is involved in microtubule binding [45], it is plausible that the CC1 domain also interacts with the heavy chain. The CC1 domain might have a secondary binding site for dynein, which may influence the microtubule-binding site of dynein, such as the stalk [46] or strut [47].

We further found that the CC1 domain affected the microtubule-binding ability of dynactin itself. The number of 1B∆CC1 mutant molecules on microtubules (8.0) was higher than that of the 1B isoform (0.6) (Fig 4c), and the microtubule-binding affinity of the 1B∆CC1 mutant was similar to that of the 1BN-GCN4 fragment as the dimerized CapGly fragment (Fig 4c). Thus, the CC1 domain may inhibit the microtubule-binding ability of the CapGly domain (Fig 5, left, red solid line). The number of 1A isoform molecules on microtubules, in contrast, was 6.6, which was much higher than that of the 1B isoform (0.6) (Fig 4c). As the 1A isoform has the K-rich domain, the K-rich domain represses the inhibitory effect of the CC1 domain (Fig 5, left, blue solid line). Furthermore, the repression by the K-rich domain of the 1A isoform might not be complete, because the binding durations of the 1A isoform (2.0 s) was still lower than that of the CapGly fragment (2.9 s) in a previous study [28]. Thus, DCTN1 has two regulatory domains, K-rich and CC1, which interact intramolecularly with each other and change the microtubule-binding affinity of dynactin.

Although the 1A isoform induced unidirectional movement of the DD complex, the number of dynein molecules on microtubules with the 1A isoform was slightly lower than that of dynein alone (Fig 2b). Moreover, we detected an increase in the fraction of stationary molecules of the DDB complex with the 1B isoform (Fig 1c). These results suggest that the properties of the two isoforms are not exclusive, and that dynactin possesses opposing properties and the distinctive function of each isoform is constituted by a predominated one of opposing properties of the regulatory domains in DCTN1 (Fig 5, left).

In a cell, the DD complex localizes at microtubule plus ends [10,16,48] or vesicles [16]. A previous *in vitro* reconstitution study only found the dynein–dynactin–EB1 complex at the microtubule plus end [49], and it is known that EB1 binds to the CapGly domain [50,51]. Our findings suggest that the CC1 domain of the 1B isoform prevents the dynein–dynactin–EB1 complex from minus end directed movement by dynein, and the complex was found only at the plus end by the aid of +TIPs. In vesicle transport, vesicles with both dynein and kinesin can exhibit plus end directional movement along microtubules [16], because the 1B isoform suppresses dynein motility. The inhibitory effect of the CC1 domain of the 1B isoform on the microtubule-binding ability of the DD complex is involved in the passive transportation toward the plus end direction. The promotion and suppression of dynein motility may be crucial for correct localization and functions of dynein within a cell.

## Acknowledgements

We thank K. Saito for helpful discussions and K. Matsuda for technical assistance. This work was supported by JSPS KAKENHI Grant Numbers 15H01310 and 16H04772 to Y.Y.T.

## Author contributions

T.K. and Y.Y.T. designed the experiments. T.K. and T.Mi. performed the experiments. T.K. and T.Mu. cloned and purified proteins. T.K. and Y.Y.T. analysed the data and wrote the manuscript.

## Supporting information

**S1 Fig. SDS-PAGE of purified dynactins.** Lane 1: marker; lane 2: 1A isoform; lane 3: 1B isoform; lane 3: 1B∆CC1 mutant.

**S2 Fig. Mean square displacement (MSD) plots.** MSD plots of 1A (•) and 1B (○) isoforms. Mean ± S.D. The diffusion coefficient (*D*) was calculated by the following formula: MSD = 2*Dt*.

**S1 Table. Quantification of dynein molecules behaviors on microtubules.**

**S1 Movie. The 1A isoform induces unidirectional movement of dynein.** Left panel: Alexa 647-labeled dynein; middle panel: GFP fused 1A isoform; right panel: merged movie. The playback speed is 10-fold faster than the acquisition speed. Scale bar is 1μm.

